# PICKER-HG: a web server using random forests for classifying human genes into categories

**DOI:** 10.1101/681460

**Authors:** Fabio Fabris, Daniel Palmer, Zoya Farooq, João Pedro de Magalhães, Alex A Freitas

## Abstract

**Motivation:** One of the main challenges faced by biologists is how to extract valuable knowledge from the data produced by high-throughput genomic experiments. Although machine learning can be used for this, in general, machine learning tools on the web were not designed for biologist users. They require users to create suitable biological datasets and often produce results that are hard to interpret.

**Objective:** Our aim is to develop a freely available web server, named PerformIng Classification and Knowledge Extraction via Rules using random forests on Human Genes (PICKER-HG), aimed at biologists looking for a straightforward application of a powerful machine learning technique (random forests) to their data.

**Results:** We have developed the first web server that, as far as we know, dynamically constructs a classification dataset, given a list of human genes with annotations entered by the user, and outputs classification rules extracted of a Random Forest model. The web server can also classify a list of genes whose class labels are unknown, potentially assisting biologists investigating the association between class labels of interest and human genes.

**Availability:** http://machine-learning-genomics.com/

## Introduction

The increasing volume of freely available biological data from sources such as the Gene Ontology (GO)[1], BioGrid [2], and GTex [3] has enabled the use of several machine learning methods to assist biologists investigating their data [4, 5] using free and open source tools such as scikit-learn^[1]^ and WEKA^[2]^.

However, as far as we know, there is no online tool that applies the standard classification workflow from the machine learning field to biological data. Our freely available web server, the “PerformIng Classification and Knowledge Extraction via Rules using random forests on Human Genes (PICKER-HG)” web server is designed to fill this niche: it is capable of reading user data in the form of class labelled human gene lists and then preparing them for classification. Depending on user preferences, the genes are then annotated with either GO terms, Protein-Protein Interactions or baseline expression levels (from GTex). The selected annotation type is then used by a sophisticated classification algorithm (random forests) to predict the class labels of the provided genes and to extract interpretable IF-THEN-ELSE rules with good predictive power directly from the classification model.

We hope that our web server will be useful to biologists exploring human genes by providing data-driven insights about various complex biological phenomena, possibly assisting in the functional classification of gene sets and the prioritization of candidate genes for further study.

## Methods

The classification task handled by this server is the computational problem of inducing a model that maps given genes to class labels (e.g. whether or not a gene is associated with ageing) using annotations describing properties of each gene (e.g. functional annotations or expression levels in different tissues).

To perform this task, usually, two sets of genes are available – a training set (which is used to build the model) and a testing set (the candidate genes). The training set contains genes for which the class label is already known, such as genes previously associated with ageing. The testing set, on the other hand, contains genes for which the class label is not known, such as genes not known to be associated with ageing.

### Data sources

To compile the training and testing datasets, one needs to describe the instances (genes) using numerical features. One of the advantages of using our server is that once the Entrez Ids. of the genes and classes are defined by the user, the training and testing datasets are automatically generated. The PICKER-HG web server uses 3 sources of data to this end: Gene Ontology (GO) terms, Protein-Protein Interactions (PPI), and GTex baseline expression levels.

GO features encode the information of which GO terms are associated with a gene (an instance). A value of ‘1’ for this feature means that the gene is known to be associated with the GO term. A value of ‘0’ means that the gene is not known to be associated with the GO term. We have used GO annotations from GO release ‘2017-03-14’.

PPI features encode for each gene, the list of proteins that interact with the products of the gene. In practice, a value ‘1’ for this feature indicates that the gene (or gene product) interacts with a given protein; a value ‘0’ indicates that there is no evidence for that interaction. We have used PPIs from BioGRID (version 3.4.146).

Lastly, GTex features encode the expression value of the gene across several tissues. For more information about the types of expression values, please read the “Help” section of our web server. We have used GTex version ‘2016-01-15 v7’.

### Training the Random Forest

Before applying the classification model to the testing set it should first be validated. We use the popular 10-fold cross-validation procedure to estimate the error of the classifier.

Once the model has been validated, if its estimated predictive performance is satisfactory, it can then be used to classify the genes in the testing set and to extract potentially useful information about the underlying classification problem. In this work we use a popular classification algorithm called Random Forest (RF)[6]. RFs models are formed from several Random Trees (a type of Decision Tree) that achieve good predictive performance [7, 8] and are interpretable to some extent, since they consist of interpretable Decision Trees.

However, it is not practical to interpret all trees in the forest, since there are usually many of them and their predictions are often contradictory. Instead, we propose a “finer-grained” interpretation, converting trees to classification rules and then pruning the rules to improve their generality and reduce overfitting.

### Rule extraction technique

We can extract potentially useful knowledge learnt by the RF in the form of classification rules. Note that a Random Forest is equivalent to a set of IF-THEN rules, one rule for each path from the root node of a decision tree to a leaf, where a prediction is made. Given a set of such rules extracted from the RF, which usually comprises thousands of rules, we can find a subset of rules with high predictive performance and return this rule set to the user.

Selecting a measure of predictive performance from the many available measures [9] is a subjective choice and using a single measure of predictive performance to rank the rules often puts more weight on one aspect of the classification performance than others. For this reason, the PICKER-HG server reports four measures of predictive performance (‘Coverage’, ‘Hits’, ‘Precision’, ‘*hits* − *error*’, defined next).

‘Coverage’ is the number of genes covered by the rule, i.e., the number of genes satisfying all conditions in the rule; ‘hits’ is the number of genes among the covered genes that were correctly classified by the rule; ‘precision’ is equal to ‘hits’ divided by ‘coverage’. The default ranking criterion (from highest to lowest value) is according to the formula *hits* − *errors*, in other words, the number of correctly classified genes minus the number of incorrectly classified genes.

We have used the following algorithm to extract rules from the RF: First, we take the final RF classification model and extract the equivalent rule set from it. Next, we discard all rules that satisfy at least one of the following criteria: 1) have a precision lower than 0.5, 2) cover fewer than 3 genes, or 3) have a precision smaller than the relative frequency of the class label. We call this subset of filtered rules *R*_*good*_. Next, a post-processing algorithm takes each rule in *R*_*good*_ in turn and tries to simplify it by executing two procedures, as follows.

#### Procedure 1

Search for the single condition whose removal *increases* the precision of the rule the most (if there are ties, choose an arbitrary condition). If such a condition is found, permanently remove the condition from the rule and repeat Procedure 1. If no condition is found by Procedure 1 (there is no condition whose removal increases the rule’s precision), execute Procedure 2. Note that the removal of conditions by Procedure 1 maintains or increases the precision of the rule and at the same time, maintains or increases its coverage.

#### Procedure 2

Search for the single condition whose removal increases the number of ‘hits’ the most, accepting a decrease in precision if the new precision is both greater than 0.75 and also greater than the relative frequency of the class label predicted by the rule (if there are ties, choose an arbitrary condition). If no such condition is found, return the current rule as the ‘simplified rule’; otherwise, remove such condition from the rule and run Procedure 1 again. In practical terms, executing Procedure 2 means that we are willing to accept rules with less precision if they are simpler (shorter), classify more instances correctly and still have high precision.

### Predictive performance estimation

An essential step in the classification workflow is the predictive *performance estimation* of the classification model being built. One should only interpret the results or use the predictions of a model with high predictive performance. In our web server, we use the Area Under the Receiver Operating Characteristic curve (AUROC) [9] measure. To estimate the AUROC of a model we use the 10-fold cross-validation procedure [9].

### Training the final RF model and getting the predictions for the testing instances

After the performance is estimated using the 10-fold cross-validation method and the performance of the classification model is considered adequate, we train the final classification model using all available instances, maximizing the model’s statistical support. Once this model is trained, we can use it to classify the ‘testing instances’ provided by the user, if any, and extract the rules from the RF.

### Server’s configuration

The PICKER-HG server was implemented using Java 8 Servlets and deployed in a Tomcat 8 web server running on a Ubuntu 12 operational system. The machine has 4Gb of RAM memory and an Intel Xeon E5-2680 processor.

## Results of an use case

The home page of the web server can be seen in the supplementary figure “home.png”. The figure shows on the left-hand side the functionalities of the server, grouped by steps, as follows.

- Step 1
  - Load dataset
  - Dataset statistics
- Step 2:
  - Train model.
- Step 3:
  - See predictive accuracy and testing results.
  - See If-Then rules.
  - Download the results.

Each step will be explained next.

### Step 1: Loading user data

After clicking on the ‘Load dataset’ button the interface shown in Figure 1 is displayed.

**Figure 1.**
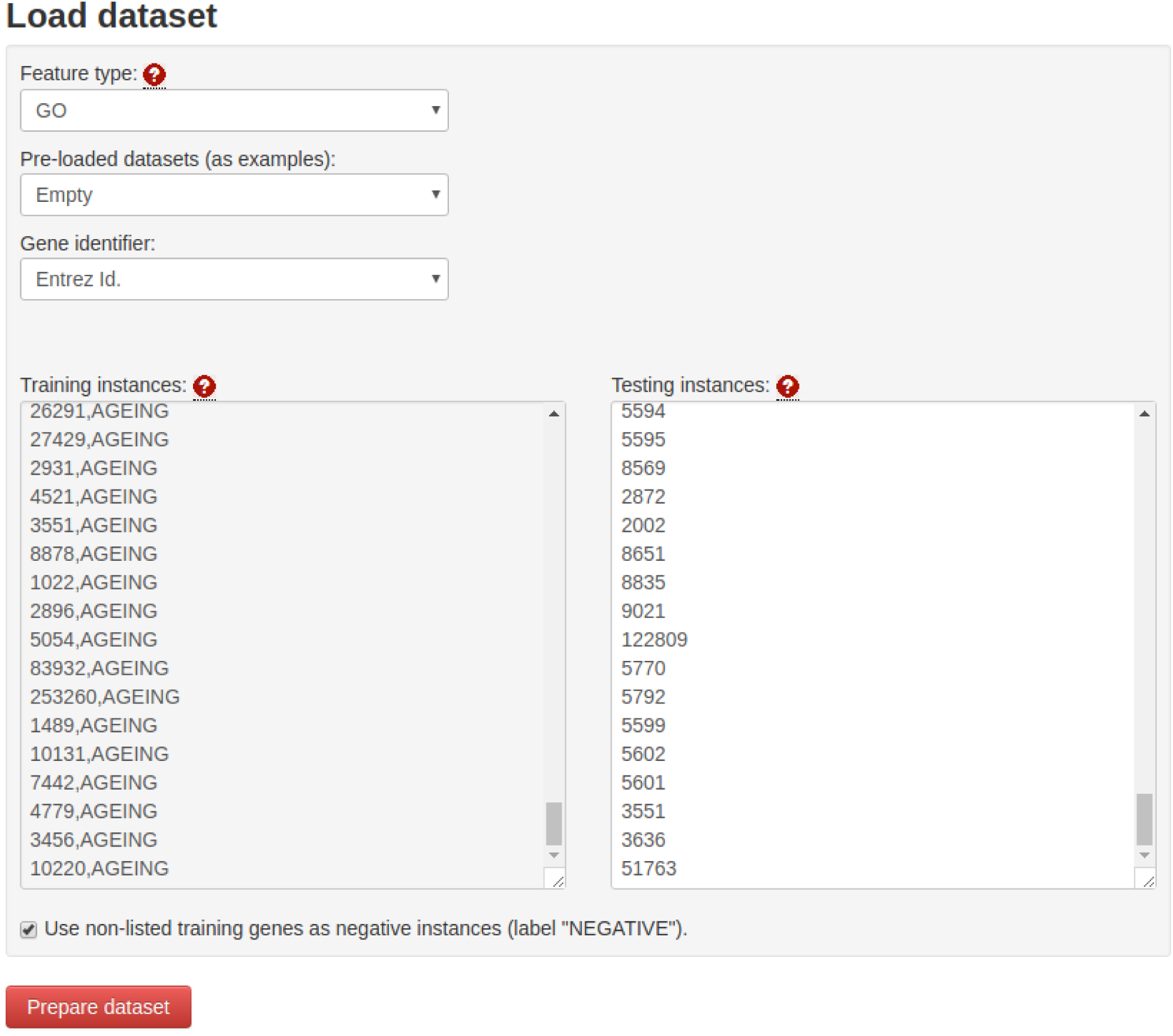
The load dataset panel is used to load the user’s data into the web server. In this panel the user can select the feature type used by the web Random Forest (RF) algorithm, use one of a list of pre-loaded datasets (as examples), select which gene identifier the user wishes to use, enter the list of training instances (gene ids) and the list of testing instances. If the user ticks the box at the end of the page, all genes not present in the training list will be labelled as the ‘NEGATIVE’ class and used as such during the training of the RF algorithm.

Before loading the gene list, the user must select a gene representation by using the selector titled ‘Feature type’. In our web server, the instances to be classified are human genes and the feature types are Gene Ontology (GO) terms [1], Protein-Protein-Interactions (PPIs) [2], or baseline expression GTex features [3]. For a given gene, a GO term (or PPI) feature with value ‘1’ encodes the information that the GO term (or PPI) is known to be associated with the gene, whereas a value ‘0’ means that the GO term (or PPI) is not known to be associated with the gene. The baseline expression GTex features encode the expression levels of the gene across several tissues in terms of absolute expression in RPKM (Reads per Kilobase Million), gene expression rank across tissues and relative expression across tissues.

The user also needs to provide a list of pre-labelled human genes (using the text box labelled ‘Training instances’) and optionally, a list of testing genes (using the text box labelled ‘Testing Instances’) with no label information – the testing genes’ labels will be predicted.

As a case study, suppose that we are interested in analysing human genes associated with ‘ageing’ according to the GenAge [10] database (supplementary file ‘ageing genes.csv’). In our case study, these genes will be used as inputs for the PICKER-HG web server, with the objective of uncovering potentially biologically relevant rules that “explain” how genes are classified into “ageing-related” (positive class) or “non-ageing-related” (negative class). Hence, the 307 genes retrieved from GenAge are considered instances of the positive class, and we set all human genes not in the list as ‘Negative’ by selecting the option ‘Use non-listed training genes as negative instances.’.

To choose the ‘testing genes’, let us consider the 137 human genes in the ‘Insulin signalling pathway’ (retrieved from KEGG version 88.2). Note that these genes will be automatically removed from the set of ‘Negative’ genes that were automatically added to the *training* instances.

To keep this study case simple, let us use the ‘GO’ feature type, as shown in Figure 1.

After all configurations are entered into the web server and the user clicks on the ‘Prepare dataset’ button, the dataset will be loaded into the system and the user will be able to go to step 2: ‘Train model’.

### Step 2: Train the model

In this step, the user must click the button ‘Train model’ and then click on the ‘Start training’ button to start the training of the RF. The progress of the various phases of the RF training is shown in the right-hand side panel.

### Step 3, Part one: Estimating predictive performance and finding targets for further biological evaluation

By clicking on the ‘See predictive accuracy and testing results’ button the user will be able to analyse the predictive performance and predictions of the web server.

To save space, only the top-5 testing human genes in the ‘Insulin signalling pathway’, in terms of the probability of being associated with ageing according to the RF model, are presented in Figure 2. This figure shows a list of ‘testing’ genes ordered by the aforementioned probability.

**Figure 2.**
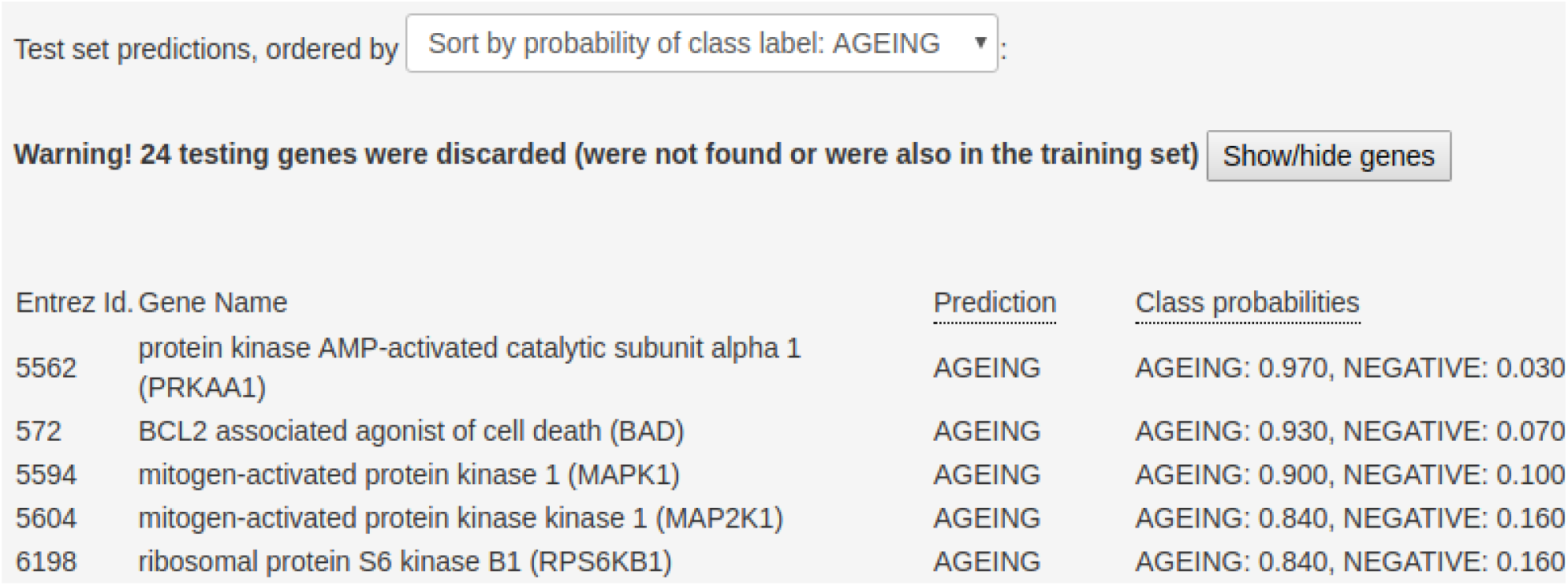
List of top-5 genes ordered by the probability of being associated with the ‘ageing’ class, according to the RF model.

A biologist could use this list to narrow down the list of genes, selecting fewer genes for further research.

In addition to the possible targets, the web server outputs the predictive performance of the RF model, estimated using a 10-fold cross-validation procedure. In our current example, we achieved an AUROC of 0.8937, which can be considered high for biological datasets.

### Step 3, Part two: Extracting the rules

Next, by clicking on the ‘See If-Then rules’ button, the user can analyse the simplified rules extracted from the RF model after running the procedures described in the section ‘Rule extraction technique’. We highlight one of the rules in Figure 3.

**Figure 3.**
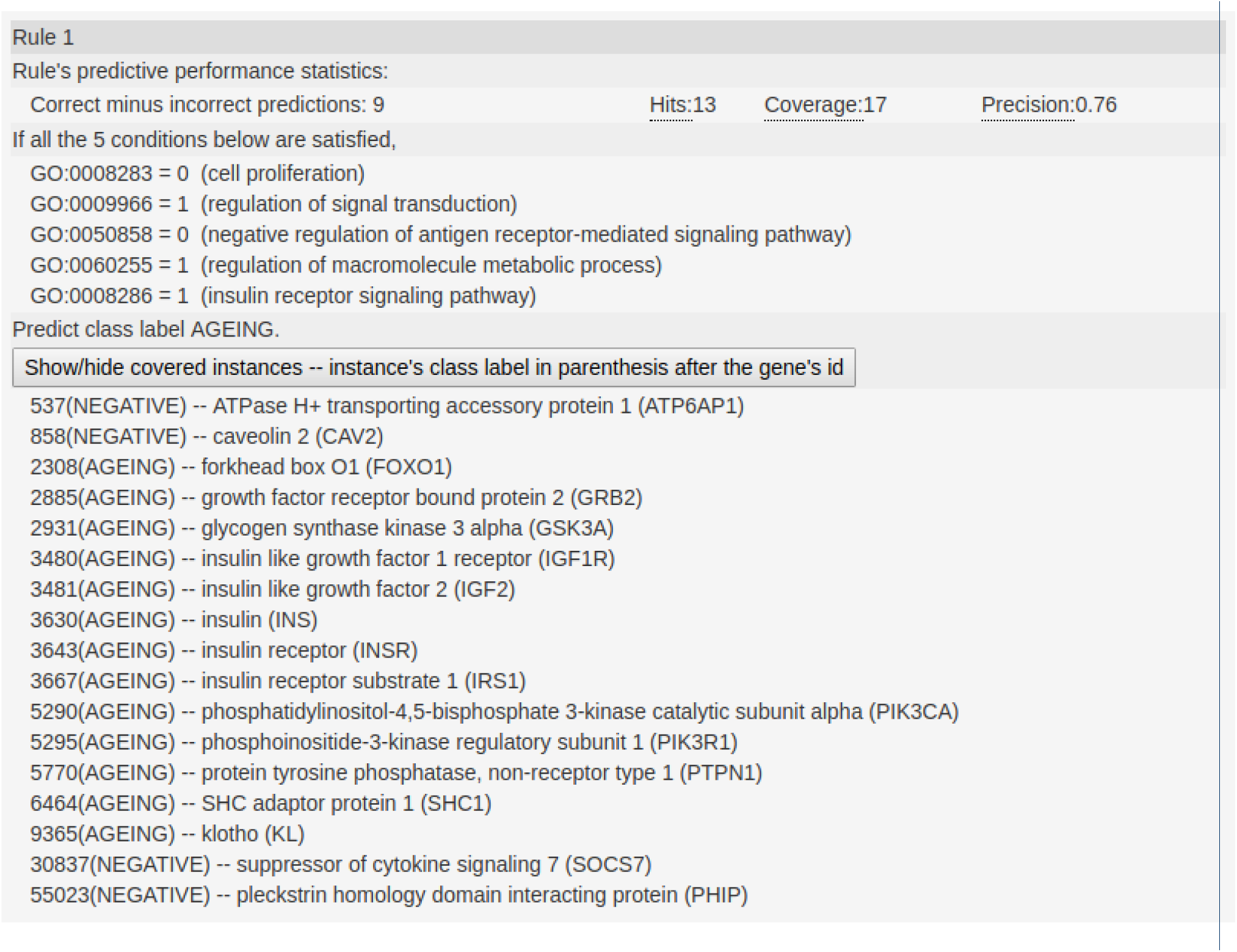
A classification rule automatically generated by the system classifying instances with the class label ‘AGEING’. The first line of the rule contains the class label being predicted (AGEING). The third line contains statistics about the predictive performance of the rule. Lines 5 to 9 contain the conditions that must be satisfied in order for an instance to be labelled as ‘AGEING’ by this rule. Finally, the last lines display the training instances (genes) captured by this rule and their actual class label, according to the data supplied by the user.

This rule shows that 17 training genes satisfy all the following conditions: they are associated with regulation of signal transduction, regulation of macromolecule metabolic process and insulin receptor signaling pathway, and at the same time, are not known to be related with cell proliferation and negative regulation of antigen receptor-mediated signaling pathway. This rule may give some important biological insights on connections between the predicted class label (‘ageing-related’) and the GO terms that were included in the “IF part” of the rule.

## Conclusion

Our web server is a flexible machine learning tool, being able to automatically create a dataset for human gene classification (given only a list of gene ids), and then training and evaluating a Random Forest model (a powerful type of machine learning method). The web server can provide valuable information in terms of retrieving testing set predictions and highly predictive rules from the Random Forest models.

## Limitations

Our web server currently only supports human genes and three feature types (GO, PPI and baseline expression from GTex), but only one feature type can be chosen when running the Random Forest algorithm. Also, we only support the AUROC measure of predictive performance. As future work, we plan to include more feature types, add a functionality to use more than one feature type at once and to support more measures of predictive performance (besides the AUROC) for the whole model.

## Supporting information

Target genes

Ageing-related genes

Home page

## Declarations

### Ethics approval and consent to participate

Not applicable.

### Consent for publication

Not applicable.

### Availability of data and materials

Link to web server: http://machine-learning-genomics.com/

### Author’s contributions

FF implemented the web server and wrote the manuscript. All other authors reviewed the manuscript and the web server. All authors read and approved the final manuscript.

## Acknowledgement

Roberto Avelar, from the University of Liverpool’s Intergrative Genomics of Ageing Group, helped test the web server.

## Footnotes

[1] https://scikit-learn.org/

[2] https://www.cs.waikato.ac.nz/ml/weka/

