## Supplementary material for "PICKER-HG: a web server using random forests for classifying human genes into categories": Home page

STEP 1

Load dataset

Dataset statistics

STEP 2

Train model

STEP 3

See predictive accuracy and testing results

See If-Then rules

Download the results

Home
Help!

### RFCRE Web Server

**(Random Forest Web Server for Classification and Rule Extraction of Human Genes)**

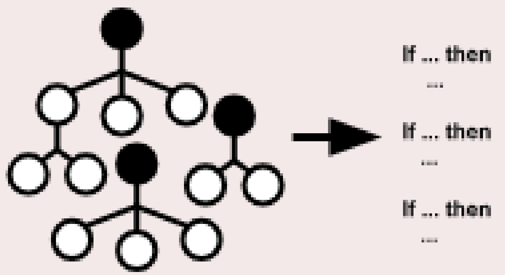


RFCRE is an easy-to-use service so that biologists can apply a state-of-the-art data mining tool to their data.

### Home

#### Quick start guide

- Step 1: Click on "Load dataset" to load your training genes and testing genes. Optionally, after you load your data, you may check statistics about your dataset by clicking on "Dataset statistics"
- Step 2: Click on "Train model" to train the classification model, this may take some time.
- Step 3: Check the testing set predictions and predictive accuracy estimation by clicking on "See predictive accuracy and testing results". To see the rules generated by the system, click on "See If-Then rules".

#### What this server can do:

- Output predictions: Estimate the probability non-labelled genes belonging to the classes defined by the user, assisting biologists identifying possible targets for further analysis.
- Generate rules: Automatically find rules that "explain" the classification of the system, possibly giving new biological insights to the users.

For more detailed information, please continue reading below.
